# Invasive plant soil legacies shape microbial function and community organization under short-term carbon and nitrogen amendments

**DOI:** 10.64898/2026.03.29.715081

**Authors:** Sawsan Hless, Aseel Sadeq, Maya Ofek-Lalzar, Yoni Gavish, Maor Matzrafi, Keren Yanuka-Golub

## Abstract

Plant invasion can modify soil microbial communities and ecosystem processes through plant-soil feedbacks, yet it remains unclear whether these effects are expressed mainly through taxonomic turnover or through shifts in microbial function and interaction structure. We tested how soil legacy generated by the invasive *Conyza bonariensis*, the native *Helminthotheca echioides*, or unplanted control soil influenced short-term microbial responses to standardized amendments and plant-derived inputs. In Experiment 1, conditioned soils were amended with water, cellulose, or ammonium and analyzed for extracellular enzyme activity, qPCR-based gene abundance, bacterial community composition, and family-level co-occurrence networks. In Experiment 2, the same soil legacies were exposed to water, glucose, or sterile root exudates from native or invasive plants. Native- and invasive-conditioned soils differed significantly in composition, but they were not consistently distinguished by strong indicator taxa, indicating that legacy effects were expressed mainly through redistribution of shared taxa rather than community turnover. In contrast, functional responses were clearer: enzyme activity and nirS abundance showed strong soil-legacy dependence, and network analysis revealed that invasive-conditioned soil supported a denser, more positive, and more compact family-level association structure than native-conditioned soil. In Experiment 2, invasive root exudates produced stronger short-term functional-based differentiation among soil legacies than native exudates, especially for extracellular enzymes. Together, the two experiments show that plant invasion can leave a persistent belowground legacy that is expressed primarily through functional filtering and network rewiring of a broadly shared microbiome, rather than through major taxonomic turnover alone.

## 1. Introduction

Plant invasions are a major driver of global environmental change, exerting strong belowground impacts via plant–soil feedbacks (PSFs) that reshape microbial communities and biogeochemical cycling (Bardgett and Van Der Putten, 2014; Ehrenfeld, 2010, 2003; Van der Putten et al., 2013). Through roots and litter, invasive plants alter resource inputs and soil chemistry, shifting microbial functional potential (e.g., extracellular enzyme activity, nitrogen-transforming capacities, and carbon use), thereby leaving legacies that persist after plants are removed (Hawkes et al., 2005; Kourtev et al., 2002; Liao et al., 2008). These feedbacks can either reinforce or mitigate invasion success and have long-term implications for soil health and ecosystem stability (Suding et al., 2013).

Invasive plants can also alter the physical structure of the soil by accelerating organic-matter decomposition, altering aggregate stability, and changing infiltration and drainage (Lone et al., 2019; Unger et al., 2022; Weidenhamer and Callaway, 2010). These changes feedback on biogeochemical cycling of carbon and nitrogen, altering soil ecological dynamics (Levine et al., 2003). In parallel, studies have shown that plant invasions change the quantity and quality of soil nutrients, with downstream effects on native plant growth (Sardans et al., 2017).

Microbial functional indicators, such as extracellular enzyme activity (EEA), nitrogen-transforming gene abundance (e.g., *nirS* for denitrification), and taxonomic composition can serve as early signals of invasion-driven change (Zhou and Staver, 2019). Increases in nutrient-releasing enzymes following invasion can accelerate cycling, potentially creating nutrient-rich conditions that favor invader persistence. These microbial shifts capture both immediate functional change and longer-term soil legacy effects. To resolve these dynamics, coupling functional assays with molecular profiling is essential, providing an integrated view of how past plant activity conditions microbial capacity to respond to new carbon or nitrogen inputs.

Root exudates are a key mechanism that modulate microbial activity, reshape community composition, and prime soil organic matter decomposition (Badri and Vivanco, 2009; Canarini et al., 2019; Kuzyakov and Blagodatskaya, 2015). These shifts translate to changes in ecosystem function by altering Carbon and Nitrogen cycling and greenhouse-gas pathways (Butterbach-Bahl et al., 2013; Negesse et al., 2025; Philippot et al., 2013). While competition with native plants can further modify PSFs (Fahey and Flory, 2022), a critical outstanding question is how plant-conditioned soil legacies alone influence short-term microbial functional responses to new carbon or nitrogen inputs.

We address this question by isolating plant–soil legacy effects and testing how they shape short-term microbial functional responses to new C and N inputs, using two annual plants that belong to the Asteraceae family with contrasting invasion status: the invasive *Conyza bonariensis (L.)*, a widespread, stress-tolerant weed with high propagule pressure and reported allelopathic effects leading to the displacement of native plant species and alterations in soil microbial processes (Djurdjević et al., 2011; Zhang et al., 2020). *C. bonariensis* is considered one of the most widespread and problematic invasive weeds globally (Bajwa et al., 2016; Weaver, 2001). Native to South America, it has successfully established in numerous regions, including the Mediterranean basin. In Israel, *C. bonariensis* has expanded rapidly across both terrestrial and semi-aquatic ecosystems, including roadsides, agricultural lands, and wetland edges (Galil et al., 2021). For the native species, we have used *Helminthia echioides* (L.) Gaertn., commonly known as bristly ox-tongue, a native annual herb, widely distributed across Mediterranean and semi-arid regions of Israel (Danin and Fragman-Sapir, 2016). It is typically found in disturbed habitats, fields, and seasonally inundated areas, where it contributes to early-season vegetation cover and supports local biodiversity. Like *C. bonariensis*, it follows a winter-spring growth cycle, germinating with the first rains of the season and flowering by late spring. *H. echioides* is ecologically significant in maintaining native plant community structure and serves as a resource for pollinators and soil-dwelling fauna. Unlike its invasive counterpart, *H. echioides* tends to coexist with other species and does not display aggressive competitive behavior.

We tested whether invasion-driven soil legacy or short-term resource inputs more strongly determines microbial responses. Using soils conditioned by the invasive *Conyza bonariensis*, the native *Helminthotheca echioides*, or unplanted, we performed two short incubation experiments. In Experiment 1, soils received water, cellulose, or ammonium, and we quantified extracellular enzyme activity, gene abundance proxies, community composition, and microbial association networks. In Experiment 2, soils received water, glucose, or root exudates from native or invasive plants to test whether biologically relevant inputs elicit legacy-dependent responses. We hypothesized that plant–soil legacy would establish the baseline state of the microbiome through persistent rhizodeposit- and exudate-mediated selection, whereas short-term resource inputs would induce context-dependent shifts whose magnitude and direction depend on that preconditioned community rather than fully resetting it (Hannula et al., 2021; Hawkes and Keitt, 2015; Hu et al., 2018; Ling et al., 2022; Nannipieri et al., 2023).

## 2. Materials and methods

### 2.1 Soil source and properties

Bulk soil for the conditioning experiment was collected in the Newe Ya’ar region (northern Israel). Particle-size analysis showed 14.0% sand, 36.4% silt, and 49.6% clay. Prior to use, soil was air-dried, passed through a 2-mm sieve to remove coarse debris, and homogenized. This Mediterranean alluvial soil typically exhibits high water-holding capacity and slow drainage, factors likely to influence microbial activity and plant–soil interactions during the experiment.

### 2.2 Soil conditioning by plant identity

We established three soil-legacy treatments using the same base soil: Control (unplanted) soil maintained without plants; Native-conditioned soil conditioned by growth of the native *H. echioides*; and an Invasive-conditioned soil conditioned by growth of the invasive *C. bonariensis*. After conditioning, soils from each treatment were harvested and used in subsequent incubations and analyses. Seeds of each species were sown into plastic containers (19 cm * 15 cm) filled with Newe Ya’ar soil and maintained in a controlled-environment chamber at 22/27 °C (night/day), with a 14h photoperiod and 700µmol m^−2^ s^−1^ photosynthetic photon flux density (fluorescent lighting). In each container, 60 seeds were sown, upon germination, plants were grown for three more weeks. Containers were watered daily to maintain consistent soil moisture, avoiding both overly dry and waterlogged conditions.

After a conditioning period of three weeks, plants were manually removed from the containers without shaking off adhering soil, as the goal was to collect root-associated soil, i.e. soil in close proximity to the root system rather than tightly bound rhizosphere soil. The remaining soil in each container, situated near the root system was carefully collected. For each container, soil from the base of six plants was pooled to form a composite sample representing the microbial community conditioned by the specific treatment. This material represents plant-conditioned bulk soil, influenced by root exudation and plant growth, rather than rhizosphere soil. The collected soils were homogenized within each treatment group and used for baseline (Time 0) measurements, as well as for subsequent incubation experiments and analyses. To determine soil moisture content, approximately 5 g of soil were weighed and dried at 105 °C for 24 h. The resulting values were used to normalize enzymatic activity rates and gene abundance measurements across samples.

### 2.3 Experiment 1: water, cellulose, and ammonium incubations

For the first incubation experiment (Fig. 1), after plants were removed, 20 g of each soil type (native, invasive or control) were weighed into clean 100 mL Erlenmeyer flasks. Soils were then incubated with 22 mL of one of the following amendments (three replicates for each soil type): Distilled water, Cellulose solution (5 mg/mL), Ammonium chloride (NH₄Cl) solution (5.5 mg/mL). All flasks were incubated in the dark at room temperature for 44 hours, without agitation.

**Figure 1.**
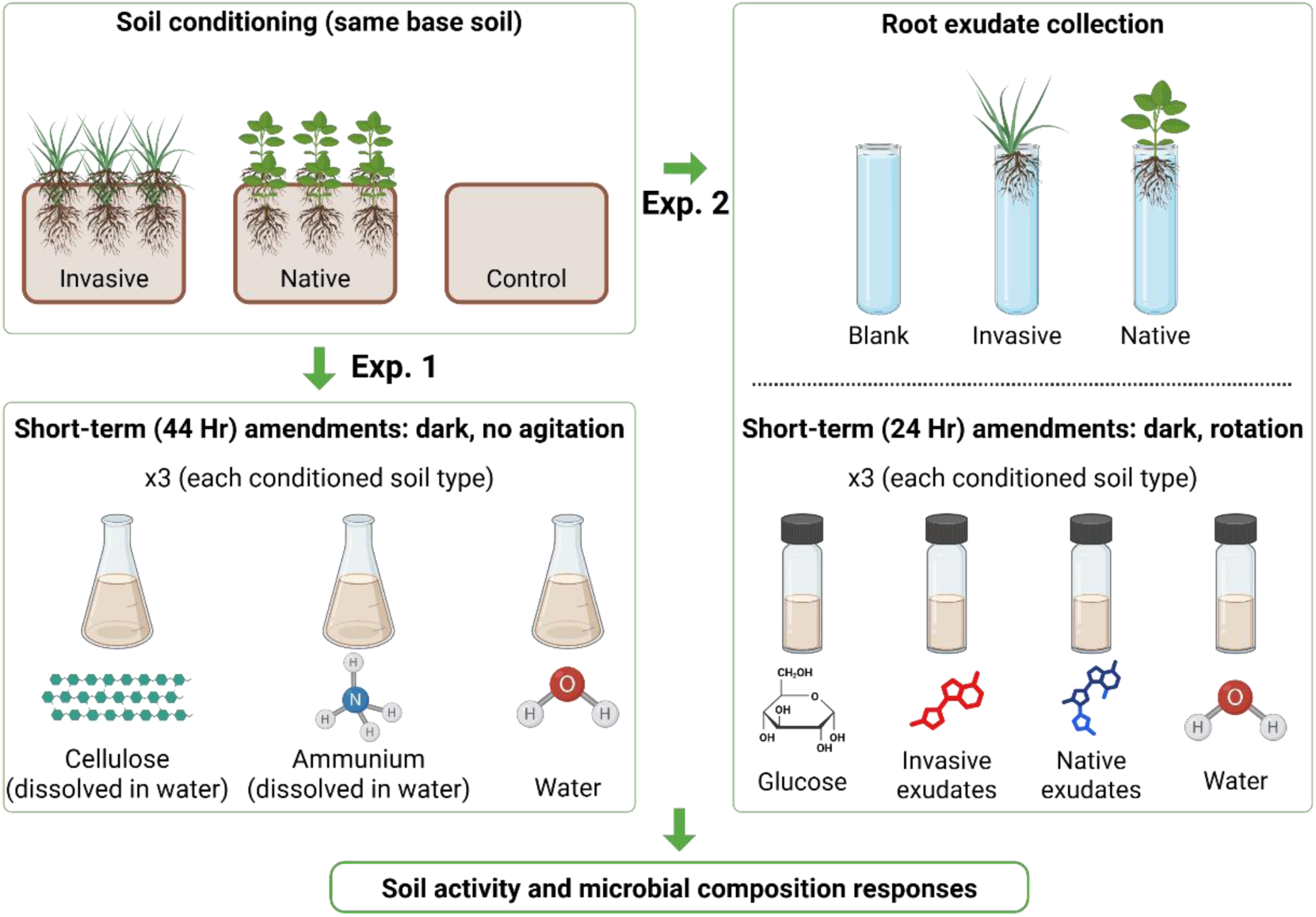
Study design and experimental workflow. Same base soil was conditioned by plant identity to generate three soils: invasive, native, and unplanted control. Experiment 1: Each conditioned soil was amended with cellulose, ammonium, or water. Experiment 2: Root exudates were collected hydroponically (blank, invasive, native) and added to each conditioned soil alongside glucose and water controls. After incubations, soils were analyzed for microbial activity and composition. Created in BioRender.

### 2.4 Root exudate collection from magenta-grown plants

Seeds of *C. bonariensis* (invasive) and *H. echioides* (native) were surface-sterilized (Supplementary information) prior to germination to minimize microbial contamination during root exudate collection. Sterilized seeds were sown in Magenta boxes containing an agar medium composed of 0.215 g L^−1^ Murashige and Skoog (MS) basal salts and 500 g L-1 Phytogel. Germination was carried out under dark conditions until sprouting (2-3 days). Once seedlings developed 2–4 true leaves, they were carefully transferred to sterile glass vials containing sterile Hoagland’s nutrient solution, as detailed in the Supplementary Information. Plants were grown under controlled conditions, and root exudates were collected weekly. After each collection, the nutrient solution was filtered through a 0.22 µm sterile membrane filter to remove particulates and potential microbial contaminants (primarily plant border cells). The filtered exudates were stored at −20 °C until use in soil incubation experiments.

### 2.5 Experiment 2: glucose and root exudate incubations

Soils conditioned by *C. bonariensis* (invasive), *H. echioides* (native), and unplanted controls were harvested after 21 days of plant growth under the conditions described above. Each conditioned soil type was then assigned to one of four amendment treatments, with three biological replicates per treatment: sterile water, root exudates collected from hydroponically grown native plants, root exudates collected from hydroponically grown invasive plants (as described above), or glucose solution as a standardized labile carbon source. For each treatment, 2 mL of the respective solution was added to 4 g dry-weight-equivalent soil placed in 60 mL amber glass vials to maintain dark conditions. The glucose treatment contained 750 µg glucose per vial. Soils were incubated at room temperature for approximately 20 h under constant mixing. Following incubation, soil slurries were centrifuged at 5,000 rpm for 5 min, and the resulting pellets were collected for downstream biological analyses. Baseline measurements were obtained from conditioned soils prior to amendment (T0) for pH, dissolved organic carbon (DOC), total dissolved nitrogen (TDN), NO_3_^−^, NH_4_^+^ DNA extraction and enzymatic activity (Fig. 1).

### 2.6 Soil chemistry

At the end of each incubation experiment, soil subsamples were centrifuged and separated into porewater and pellet fractions. Porewater were used for chemical analyses, whereas soil subsamples were used for biological assays, including extracellular enzyme activity, DNA extraction for microbial community analysis, and quantitative PCR, as described below. For porewater extraction, 20 g soil subsamples were centrifuged at 3,000 × g for 10 min, and the resulting supernatant was filtered through 0.45 µm syringe filters. Filtrates were stored at −20 °C until analysis. In Experiment 1, ammonium (NH_4_ ^+^) was quantified using a sodium salicylate colorimetric assay modified from Krom (1980). Dissolved organic carbon (DOC) and total dissolved nitrogen (TDN) were measured using a Shimadzu TOC-L analyzer (Columbia, MD, USA). Soil pH was also measured at a 1:5 soil:water ratio. In Experiment 2, soil chemical extractions were focused on the labile dissolved nutrient pool, including nitrate and ammonium. DOC and TDN were extracted with 0.5 M K_2_SO_4_ and analyzed as described above. Nitrate (NO_3_ ^−^) and ammonium (NH_4_ ^+^) were extracted and quantified according to Standard Methods 4500-NO_3_ ^−^ E and 4500-NH -F, respectively.

### 2.7 Extracellular enzyme assays

Potential activities of two hydrolytic enzymes were measured fluorometrically to assess microbial capacity for carbohydrate and nitrogenous organic-matter degradation: α-1,4-glucosidase (AG; EC 3.2.1.20) and β-1,4-N-acetylglucosaminidase (NAGase; EC 3.2.1.52). Assays were performed using 4-methylumbelliferyl-linked substrates following a modified protocol based on German et al. (2011) and Hless and Yanuka-Golub (in press). Soil homogenates were prepared fresh in sterile water, incubated with substrate in the dark for 1 h at room temperature, and fluorescence of released 4-methylumbelliferone (MUF) was measured with a microplate reader. Enzyme activity was quantified from MUF standard curves with sample-specific quench correction and expressed as µmol MUF released g⁻¹ dry soil h⁻¹. Detailed protocols for homogenate preparation, control design, incubation conditions, fluorescence standardization and measurement, and enzyme activity calculations are described in the Supplementary Methods and elsewhere (Hless and Yanuka-Golub, in press).

### 2.8 DNA extraction, PCR amplification, and amplicon sequencing

Whole-community genomic DNA was extracted from soil subsamples using the FastDNA™ SPIN Kit for Soil and quantified by NanoDrop spectrophotometry. Full-length bacterial 16S rRNA gene amplicons were generated using the primers 27F/1492R in a two-stage PCR workflow. In the first stage, target amplicons were produced using MyTaq HS Mix, and in the second stage, PacBio Kinnex adapter sequences and combinatorial dual indices were added using repliQa HiFi ToughMix. Amplicons were pooled, purified, and sequenced on a PacBio Revio platform. Initial second-stage amplification and indexing were performed at the Genomics and Microbiome Core Facility at Rush University, and final library preparation and sequencing were carried out at the DNA Services Facility, Roy J. Carver Biotechnology Center, University of Illinois Urbana-Champaign. Raw sequence data were deposited in the NCBI Sequence Read Archive under accession number PRJNA1441857. Full details of DNA extraction, PCR conditions, indexing, cleanup, and sequencing are provided in the Supplementary Methods.

### 2.9 Sequence processing and community data analysis

Full-length 16S rRNA gene amplicon data were processed in QIIME2 amplicon-2024.5 (Rush Research Bioinformatics Core at Rush University Medical Center). Raw reads were quality-checked with seqkit, primer sequences (27F/1492R) were removed with cutadapt, and denoising was performed with DADA2 via q2-dada2-ccs, generating amplicon sequence variants (ASVs). Taxonomy was assigned using a Naive Bayes classifier trained against the GTDB r202 reference database, and putative contaminants were identified with decontam on the basis of prevalence in reagent negative controls.

For Experiment 1, the initial table comprised 13,934 ASVs across 43 samples. After removal of three low-read samples (ContCell3, Cont2_1, and Cont2_2), and filtering of zero-abundance ASVs, the final dataset used for diversity and community analyses comprised 12,308 ASVs across 31 samples. For alpha-diversity analyses, counts were rarefied to 30,000 reads per sample using random subsampling without replacement (rrarefy, vegan). Observed richness, Shannon diversity, and Simpson diversity were calculated from the rarefied table, and effective Shannon and effective Simpson diversity were derived as Hill numbers. Differences in alpha diversity were tested using one-way aligned rank transform (ART) ANOVA for Soil (Control, Native, Invasive), Condition (T0, Water, Ammonium, Cellulose), and Plant (No/Yes), with Tukey-adjusted post hoc comparisons when appropriate. Full details of filtering, rarefaction, metric calculation, and statistical analysis are provided in the Supplementary Methods.

For community-composition analyses, the non-rarefied count table was normalized by cumulative sum scaling (CSS) using metagenomeSeq. The CSS-normalized ASV matrix contained 12,308 ASVs × 31 samples and was additionally aggregated to the genus level, yielding a matrix of 1,271 genera × 31 samples. Bray–Curtis dissimilarities were visualized by two-dimensional NMDS (metaMDS, vegan) at both the ASV and genus levels; the corresponding stress values were 0.0359 and 0.0386, respectively. PERMANOVA (adonis2, vegan) was used to test the effects of Soil, Condition, Plant, and Soil × Condition on community composition. In addition, a second genus-level NMDS and PERMANOVA analysis was performed on the subset containing only plant-conditioned soils (Native and Invasive); the stress value of that ordination was 0.1538. To further examine similarity patterns within the plant-conditioned subset, Bray–Curtis dissimilarities were calculated from CSS-normalized genus-level data and subjected to hierarchical clustering using average-linkage (UPGMA). Dendrograms were used to visualize grouping among samples, and Bray–Curtis distance matrices were displayed as heatmaps with soil legacy and incubation condition annotations. Differential abundance analyses were performed on the plant-conditioned subset using LEfSe and condition-specific ALDEx2 comparisons between native- and invasive-conditioned soils; full methodological details and parameter settings are provided in the Supplementary Methods.

### 2.10 Quantitative PCR

Quantitative PCR (qPCR) assays were performed to assess the genetic potential of the soil microbial communities, following established protocols (Borchardt et al., 2021). Reactions were conducted using the SsoAdvanced™ Universal Inhibitor-Tolerant SYBR® Green Supermix (Bio-Rad, Hercules, CA, USA) on a Bio-Rad CFX96 Real-Time PCR System, with data analyzed using CFX Manager Version 2.3. Each reaction (25 μL total volume) included 12.5 μL SYBR® Green Supermix, 0.3 μM of each primer, and approximately 10 ng of genomic DNA. All samples were run in triplicate. Results are reported as log₁₀ gene copies per mg of dry soil, calculated by averaging technical replicates and referencing standard curves. Negative controls containing DNA-free water were included in all runs as negative controls.

Archaeal 16S rRNA gene fragments (∼140 bp) were amplified using universal primers Arch 967F (5′-AATTGGCGGGGGAGCAC-3′) and Arch-1060R (5′-GGCCATGCACCWCCTCTC-3′) (Cadillo-Quiroz et al., 2006; Karlsson et al., 2012). Thermal cycling conditions were: initial denaturation at 95 °C for 5 min, followed by 40 cycles of 95 °C for 20 s, 59 °C for 20 s, and 72 °C for 30 s, with a final melt curve analysis. Standard curves were generated from tenfold serial dilutions (10⁻¹ to 10⁻⁶) of *Methanosarcina barkeri* genomic DNA (DSM 800, DSMZ), with amplification efficiency of 106.5% and R² = 0.99.

Several functional genes were initially screened using endpoint PCR (amoA, phoD, nirK, nirS, pmoA); however, nirS was the only gene consistently detected across all soil samples. Denitrifier abundance was quantified using cd3AF and R3cd primers targeting the nirS gene (Throbäck et al., 2004). The qPCR protocol consisted of an initial denaturation at 95 °C for 3 min, followed by 40 cycles of 95 °C for 45 s, 58 °C for 45 s, and 72 °C for 45 s, concluding with melt curve analysis. Standard curves for nirS were generated using tenfold serial dilutions (10⁻² to 10⁻⁹) of synthetic double-stranded DNA fragments (gBlocks, Integrated DNA Technologies, IDT) suspended in TE buffer (20 ng μL⁻¹ stock), based on the expected amplicon sequence (Han et al., 2023). The amplification efficiency was 91.3%, with R² = 0.99.

### 2.11 Diversity and network analyses

For network analyses, Experiment 1 ASV data were aggregated to the family level and normalized by cumulative sum scaling (CSS) using metagenomeSeq. Co-occurrence networks were then constructed in ggClusterNet (v2.0; Wen et al., 2025, 2022) separately for soil treatment (Control, Native, Invasive) and incubation treatment (T0, Water, Ammonium, Cellulose). Interaction networks for individual Soil × Condition combinations were not constructed because sample numbers per combination were insufficient for stable inference. Networks were inferred from Spearman correlations among the 100 most abundant families within each group, using thresholds of |r| ≥ 0.6 and P ≤ 0.05. Network structure was characterized using node-level topology, modular organization, graph-level properties, and robustness analyses, including random-removal simulations and natural connectivity. Additional details on filtering, normalization, network construction, module analysis, topological role classification, random-network comparisons, and stability calculations are provided in the Supplementary Methods.

For module-level summaries, taxa were assigned to edge-sign categories based on the sign composition of their incident edges within each network. Taxa with at least 10 total incident edges and a positive-edge fraction ≥0.70 were classified as mostly positive, whereas taxa with at least 10 total incident edges and a positive-edge fraction ≤0.30 were classified as mostly negative. Taxa not meeting either criterion, including those with fewer than 10 total incident edges, were classified as mixed. Network roles were assigned using Zi–Pi analysis, in which within-module degree z-score (z) quantifies local connectivity within a module and participation coefficient (p) quantifies connectivity across modules. Role categories therefore reflect the combined within- and among-module position of each taxon.

### 2.12 Statistical analyses

Statistical analyses were conducted in R version 4.5.1 using aligned rank transform (ART) ANOVA. For both Experiment 1 and Experiment 2, response variables, including extracellular enzyme activities and qPCR-based gene abundances, were evaluated across the three soil legacies generated during conditioning (Control, Native, Invasive) and the amendment treatments applied in each experiment. In Experiment 1, amendment treatments were water, cellulose, and ammonium, whereas in Experiment 2, amendment treatments were water, native root exudates, invasive root exudates, and glucose. Each soil × amendment combination included three biological replicates, and technical replicates were averaged before statistical analysis so that each biological replicate contributed a single value per response variable. Experiment 2 was designed to compare soil legacies within each amendment treatment, thus analyses were performed as separate one-way ART ANOVAs within each amendment, followed by Tukey-adjusted pairwise comparisons among Control, Native-conditioned, and Invasive-conditioned soils when appropriate. Differences were considered significant at p < 0.05.

## 3. Results

### 3.1 Baseline differences among conditioned soils

Before amendment, the physicochemical properties and baseline microbial attributes of the three conditioned soils used in Experiment 1 were characterized: unplanted control soil, native-conditioned soil, and invasive-conditioned soil (Table 1). All soils were found to be alkaline, with pH values ranging from 7.8 to 8.0. Total organic carbon (TOC) in soil porewater was found to be highest in the control soil (38.4 mg L⁻¹) and lower in the native- and invasive-conditioned soils (6.0 and 4.2 mg L⁻¹, respectively). Total dissolved nitrogen (TDN) was below the detection limit (< 5 mg L⁻¹) in all treatments. Because the control soil was not incubated and watered alongside the planted soils, but instead was collected in a dry state immediately before the conditioned soils were sampled, the higher TOC measured in the control soil was likely influenced by this difference in handling and moisture regime.

**Table 1.** Main physicochemical characteristics and baseline microbial attributes of unamended conditioned soils used in Experiment 1.

| Soil type<br>Incubation<br>Exp. 1<sup>i</sup> | pH | Total Organic<br>Carbon<br>(TOC)<br>mg L<sup>-1</sup> | Total<br>Dissolved<br>Nitrogen<br>(TDN)<br>mg L<sup>-1</sup> | NAGase<br>activity<br>$\mu\text{mol g}^{-1} \text{hr}^{-1}$<br>$p < 0.05^{\text{ii}}$ | nirS copies<br>($\times 10^2 \text{ mg}^{-1}$<br>dry soil)<br>$p > 0.05^{\text{iii}}$ | Archaeal 16S<br>rRNA gene<br>copies ($\times 10^6$<br>$\text{mg}^{-1}$ dry soil)<br>$p > 0.05^{\text{iii}}$ |
| --- | --- | --- | --- | --- | --- | --- |
| **Control** | 7.8 | 38.4 | <5 | $7.97 \pm 0.31$ | $2.12 \pm 0.20$ | $8.56 \pm 0.99$ |
| **Native** | 8.0 | 6.0 | <5 | $19.66 \pm 1.21$ | $3.76 \pm 0.68$ | $8.55 \pm 0.38$ |
| **Invasive** | 7.8 | 4.2 | <5 | $12.42 \pm 0.81$ | $3.97 \pm 0.72$ | $5.70 \pm 0.13$ |

<sup>i</sup> Values are mean ± SD of biological replicates ( $n = 3$ for NAGase; $n = 2$ for qPCR assays). For qPCR assays, technical replicates were averaged within each biological replicate before analysis. The unplanted control soil was not incubated and regularly watered alongside the planted soils during the conditioning phase. Instead, it was collected in a dry state immediately before the native- and invasive-conditioned soils were sampled. This difference in handling and moisture regime should be considered when comparing baseline soil properties, particularly TOC and enzymatic activity.
<sup>ii</sup> Between-soil differences were tested with ART-ANOVA followed by Tukey-adjusted post hoc tests on aligned estimated marginal means ( $\alpha = 0.05$ )
<sup>iii</sup> Between-soil differences were tested with ART-ANOVA, followed by Tukey-adjusted post hoc tests on aligned estimated marginal means ( $\alpha = 0.05$ ).

A set of functional and taxonomic marker genes was first screened by endpoint PCR, including mcrA, pmoA, nirS, nirK, amoA, and archaeal 16S rRNA. Of these targets, only nirS and archaeal 16S rRNA were consistently amplified across samples; the remaining markers were not detected and were therefore excluded from subsequent analyses. Baseline qPCR measurements showed numerical differences among soils, but no statistically significant differences were detected. nirS copy numbers were found to be higher in the native- and invasive-conditioned soils than in the control soil, but no significant effect of soil type was detected (ART-ANOVA, p = 0.130). Archaeal 16S rRNA gene abundance also did not differ significantly among soils (ART-ANOVA, p = 0.176), although lower mean values were observed in the invasive-conditioned soil (Table 1). In contrast, baseline NAGase activity was found to differ significantly among soil legacies before any amendment was applied (ART-ANOVA, p = 0.001). Based on Tukey-adjusted post hoc comparisons, NAGase activity was found to be highest in the native-conditioned soil, intermediate in the invasive-conditioned soil, and lowest in the unplanted control. Thus, the clearest baseline difference among conditioned soils was observed in extracellular enzyme activity rather than in the abundance of the two quantified gene markers. Because the control soil was not incubated and watered alongside the planted soils during the conditioning phase, the observed differences in enzymatic activity and TOC cannot be attributed solely to plant conditioning. Nevertheless, baseline differences associated with prior plant conditioning were detected, particularly in enzymatic potential, before short-term substrate addition.

### 3.2 Functional and gene-abundance responses to short-term amendments (Experiment 1)

In Experiment 1, β-1,4-N-acetylglucosaminidase activity, archaeal 16S gene abundance, and nirS gene abundance were measured in control, native-conditioned, and invasive-conditioned soils after amendment with water, cellulose, or ammonium (Fig. 2). Overall, the magnitude and direction of soil-type differences were found to depend on the response variable and amendment identity.

**Figure 2.**
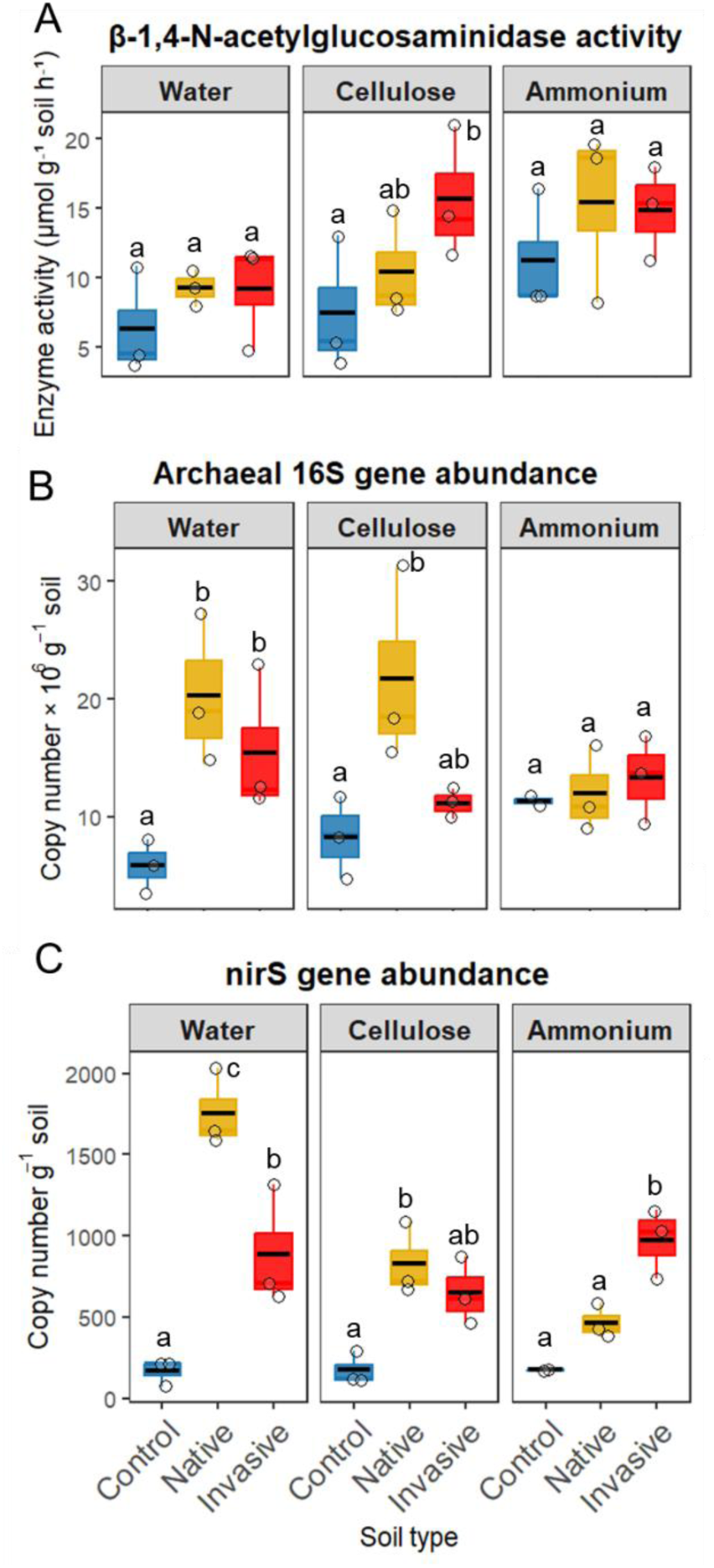
Short-term amendments differentially affected functional and gene-abundance responses of conditioned soils in Experiment 1. Responses of (A) β-1,4-N-acetylglucosaminidase activity, (B) archaeal 16S gene abundance, and (C) nirS gene abundance are shown for control, native-conditioned, and invasive-conditioned soils after amendment with water, cellulose, or ammonium. Points represent biological replicates, boxplots summarize replicate distributions, and black horizontal lines indicate means. Different letters indicate significant differences among soil types within each amendment based on ART-ANOVA followed by Tukey-adjusted post hoc comparisons (α = 0.05).

For β-1,4-N-acetylglucosaminidase activity (Fig. 2A), a significant main effect of treatment was detected (ART-ANOVA, F = 3.85, p = 0.041), whereas the effect of soil type was marginal (F = 2.66, p = 0.098), and no soil type × treatment interaction was detected (F = 0.41, p = 0.797). Averaged across soil types, activity was higher under ammonium than under water amendment (p = 0.0327). Within individual treatments, no significant differences among soil types were detected under water, whereas under cellulose a significant soil-type effect was detected (F = 8.13, p = 0.002), driven by higher activity in invasive-conditioned soil than in control soil (p = 0.0014), with native-conditioned soil showing intermediate values. Under ammonium, only a marginal soil-type effect was detected (F = 2.72, p = 0.086).

For archaeal 16S gene abundance (Fig. 2B), a strong main effect of soil type was detected (ART-ANOVA, F = 15.43, p = 0.00015), whereas no treatment effect was detected (F = 0.10, p = 0.904), and the interaction was marginal (F = 2.50, p = 0.082). Across treatments, archaeal 16S abundance was higher in native-conditioned soil than in control soil (p = 0.0001), and higher in invasive-conditioned soil than in control soil (p = 0.0087), whereas native- and invasive-conditioned soils did not differ significantly. Within treatments, significant soil-type effects were detected under water and cellulose (both F = 8.74, p = 0.0167), with native-conditioned soil exceeding control soil under both amendments (p = 0.0152), whereas no significant soil-type effect was detected under ammonium (F = 0.28, p = 0.770).

For nirS gene abundance (Fig. 2C), significant effects of soil type (ART-ANOVA, F = 19.42, p < 0.001), treatment (F = 10.21, p = 0.0012), and their interaction (F = 9.88, p = 0.00025) were detected. Because a significant interaction was observed, comparisons were interpreted within each treatment. Under water, a significant soil-type effect was detected (F = 27.0, p = 0.001), with abundance ranked as native-conditioned > invasive-conditioned > control, and all pairwise contrasts being significant. Under cellulose, a significant soil-type effect was also detected (F = 8.74, p = 0.0167), driven by higher abundance in native-conditioned soil than in control soil (p = 0.0152), whereas invasive-conditioned soil was intermediate. Under ammonium, a significant soil-type effect was again detected (F = 20.83, p = 0.0038), but the pattern shifted, with nirS abundance highest in invasive-conditioned soil that differed significantly from both control (p = 0.0033) and native-conditioned soil (p = 0.0264). Together, these results indicate that short-term amendments modified microbial responses, but that soil legacy remained an important determinant of both functional and gene-abundance patterns.

### 3.3 Community diversity and composition responses to short-term amendments (Experiment 1)

Community diversity in Experiment 1 was assessed from the filtered and rarefied ASV table using observed richness, effective Shannon diversity, and effective Simpson diversity (Fig. 3). When all soils were included, no effect of soil type was detected for observed richness, and only a marginal effect was detected for effective Shannon diversity (Fig. 3A). In contrast, effective Simpson diversity differed significantly among soils, with lower values in control soil than in both native- and invasive-conditioned soils, whereas the two plant-conditioned soils did not differ from one another (Fig. 3A).

**Figure 3.**
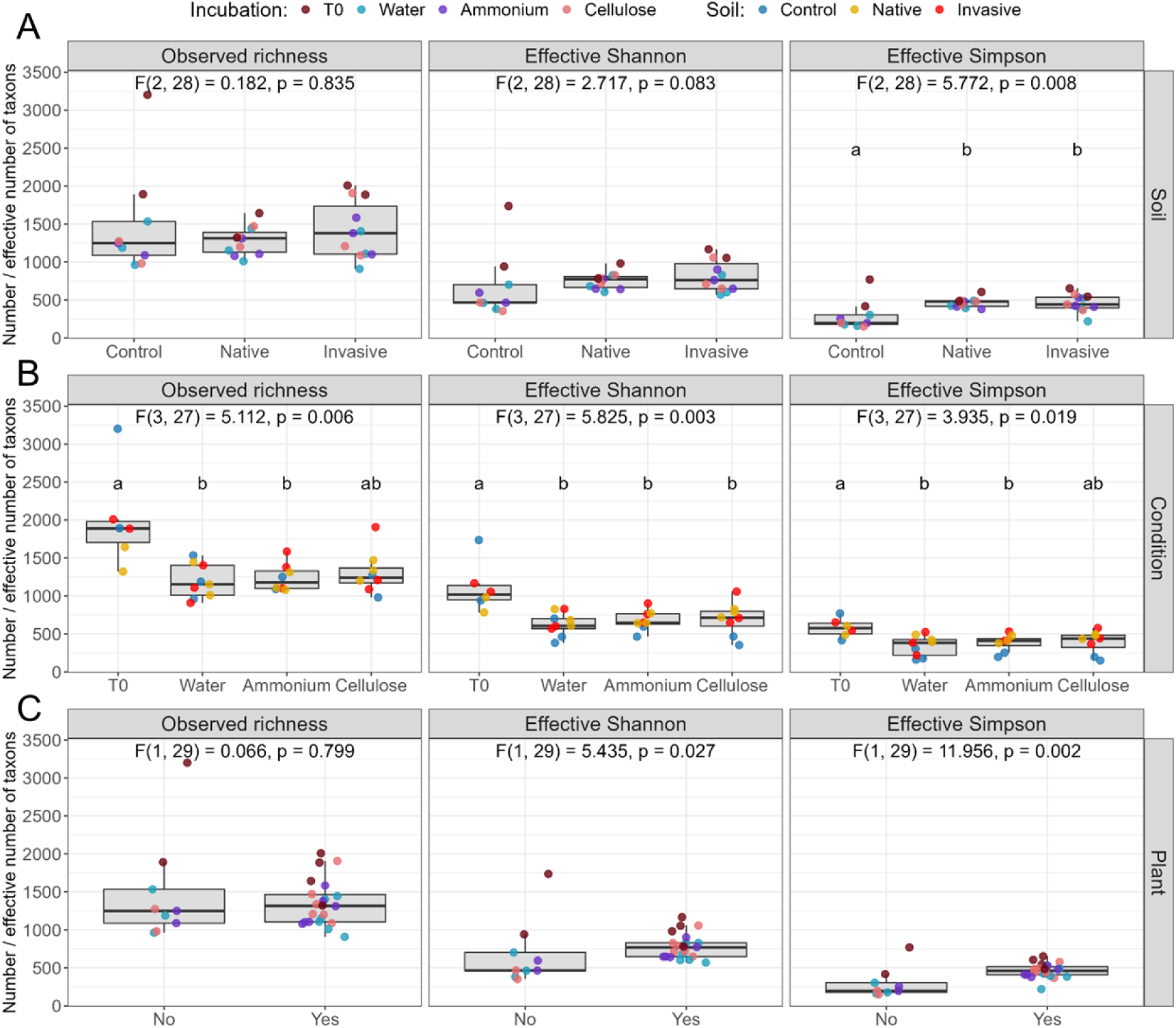
Alpha-diversity responses of microbial communities in Experiment 1. (A) Soil-type effects, (B) incubation-condition effects, and (C) plant-conditioning effects on observed richness, effective Shannon diversity, and effective Simpson diversity calculated from the filtered, rarefied ASV table. T0 represents the initial soil samples collected before incubation and amendment. Boxplots show medians and interquartile ranges; points represent individual samples. Different letters indicate significant differences among groups based on ART-ANOVA with Tukey-adjusted post hoc tests (P < 0.05).

A significant effect of incubation condition was detected for all three diversity metrics in the full dataset (Fig. 3B). Observed richness was higher at the initial sampling point collected before incubation and amendment (T0) than under water and ammonium, whereas cellulose showed intermediate values. Effective Shannon diversity was also higher at T0 than under all incubation treatments. For effective Simpson diversity, T0 exceeded water and ammonium, whereas cellulose again was intermediate (Fig. 3B). No effect of plant presence was detected for observed richness, but effective Shannon diversity and effective Simpson diversity were higher in plant-conditioned than in unplanted control soils (Fig. 3C). When the analysis was restricted to plant-conditioned soils, no effect of soil type was detected for observed richness, effective Shannon diversity, or effective Simpson diversity, indicating that alpha diversity did not differ between native- and invasive-conditioned soils. In contrast, incubation condition remained important within this subset. Observed richness responded only marginally to condition, whereas significant condition effects were detected for effective Shannon diversity and effective Simpson diversity. In both cases, diversity was highest at T0 and lowest under water, whereas ammonium and cellulose were intermediate and did not differ significantly from either group. Overall, these results indicate that community diversity in Experiment 1 was influenced more strongly by incubation condition than by differences between native- and invasive-conditioned soils, while the clearest soil-history contrast was observed between plant-conditioned and unplanted control soils.

Community-composition patterns were further examined by NMDS ordination of Bray–Curtis dissimilarities based on CSS-normalized data (Fig. 4). In the full dataset, the genus-level NMDS ordination showed that control samples were clearly separated from plant-conditioned soils, whereas native- and invasive-conditioned soils showed partial overlap (Fig. 4A). PERMANOVA confirmed significant effects of Soil (R² = 0.449, F = 41.59, p = 0.001), Condition (R² = 0.229, F = 14.15, p = 0.001), and Soil × Condition (R² = 0.219, F = 6.75, p = 0.001). Pairwise comparisons further showed that all soil contrasts were significant, including invasive vs. native. Among conditions, water, ammonium, and cellulose each differed from T0, whereas water vs. ammonium and water vs. cellulose were not significant and ammonium vs. cellulose was marginal.

**Figure 4.**
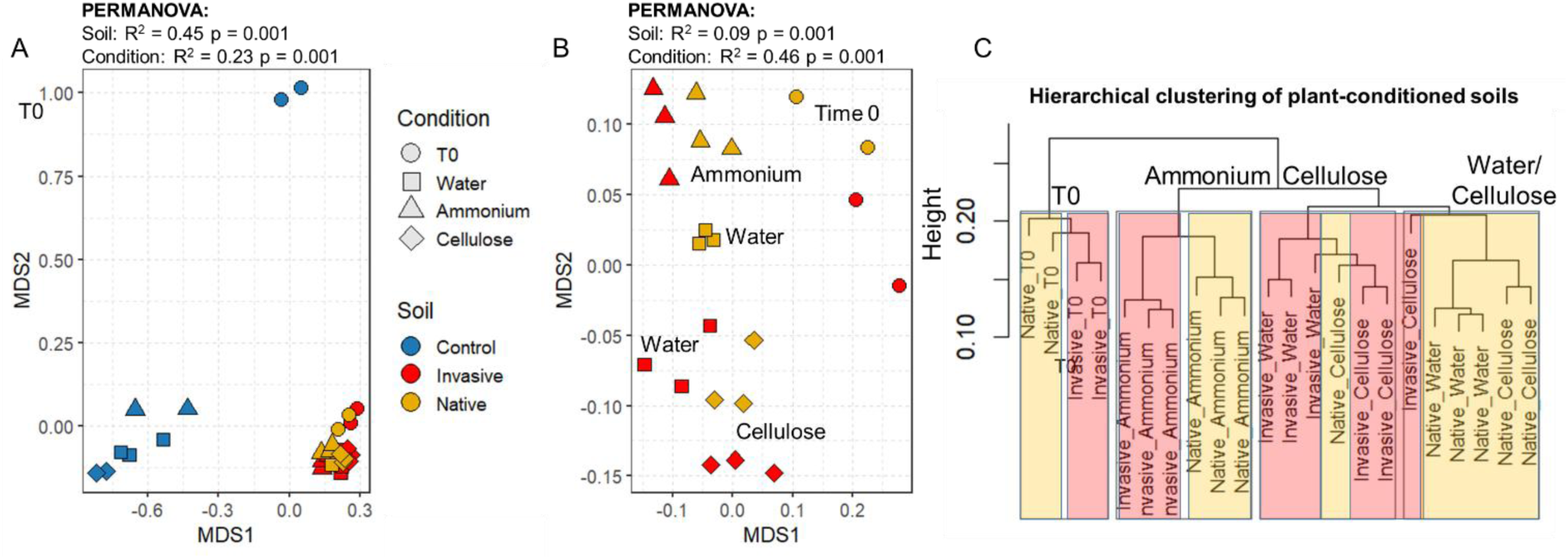
Genus-level microbial community composition in Experiment 1 based on CSS-normalized data. **(A)** NMDS ordination of all soils (Control, Native, and Invasive). **(B)** NMDS ordination restricted to plant-conditioned soils (Native and Invasive). **(C)** Hierarchical (UPGMA of the Bray–Curtis distance matrix) clustering of plant-conditioned soils based on Bray–Curtis dissimilarities. NMDS ordinations were generated from Bray–Curtis dissimilarities using two-dimensional non-metric multidimensional scaling (NMDS; *k* = 2, try = 100). Points are colored by soil type and shaped by incubation condition; T0 indicates samples collected before incubation and amendment. Stress values were 0.039 in panel A and 0.154 in panel B.

When the ordination was restricted to plant-conditioned soils, the genus-level NMDS stress increased to 0.1538 (Fig. 4B). Across plant-conditioned soils, community clustering was driven mainly by incubation condition (Fig. 4C), with native- and invasive-conditioned soils showing a weaker but significant secondary separation. This was consistent with PERMANOVA results in which condition explained most of the variance (R² = 0.4565, F = 6.72, p = 0.001), followed by smaller soil-legacy (R² = 0.0937, F = 4.14, p = 0.001) and interaction effects (R² = 0.1329, F = 1.96, p = 0.013). This pattern was evident both in the NMDS ordination and in the hierarchical clustering analysis (Fig. 4B,C). Pairwise tests within the plant-only subset showed that water differed from ammonium and T0, ammonium differed from cellulose and T0, and cellulose differed from T0, whereas water vs. cellulose remained only marginal (p = 0.041). Although PERMANOVA detected a significant separation between native- and invasive-conditioned soils, differential-abundance analyses (ALDEx2 and LEfSe) did not identify strong family-level taxa consistently distinguishing the two legacies across treatments (Supplementary Table S1-S4). Together, these results indicate that the difference between native- and invasive-conditioned soils was not characterized by a major replacement of dominant taxa. Rather, invasion legacy appeared to reshape the organization of an otherwise largely shared microbial assemblage, suggesting that the two communities may differ more in the organization and relative distribution of shared taxa than in the presence or absence of unique indicator families. To examine this further, the distribution of selected core and differential taxa was summarized first (Fig. 5), followed by analyses of family-level co-occurrence structure and network robustness (Fig. 6).

**Figure 5.**
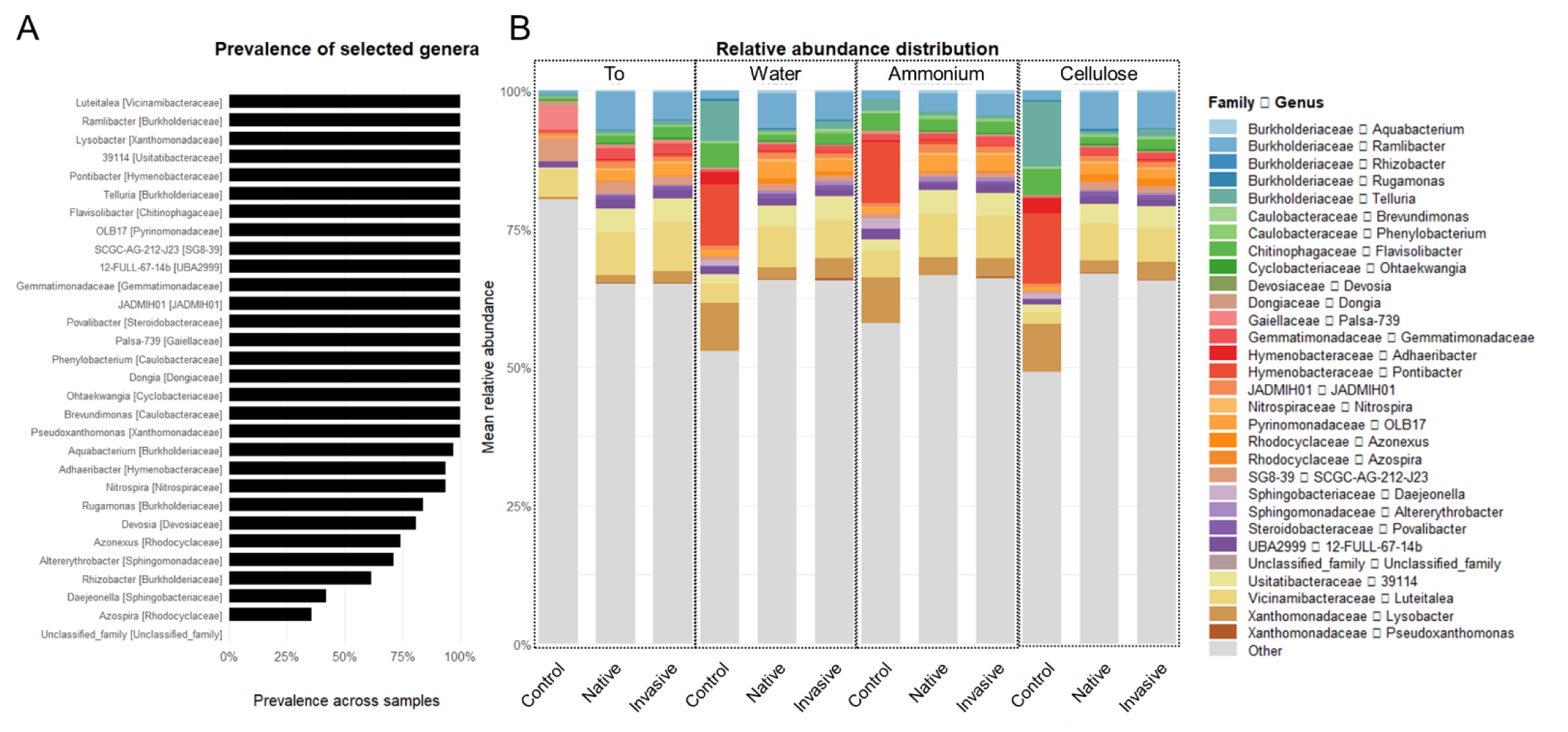
Prevalence and relative abundance distribution of selected genera in Experiment 1. (A) Prevalence of selected genera across all samples. Selected taxa were compiled from the core set together with genera identified by LEfSe across all soils and by ALDEx2 in native-versus-invasive comparisons. Family assignment is shown in brackets. (B) Mean relative abundance of the same selected genera across soil types and incubation conditions. The selected taxa represented only a subset of the total community and the remaining fraction is shown as Other.

**Figure 6.**
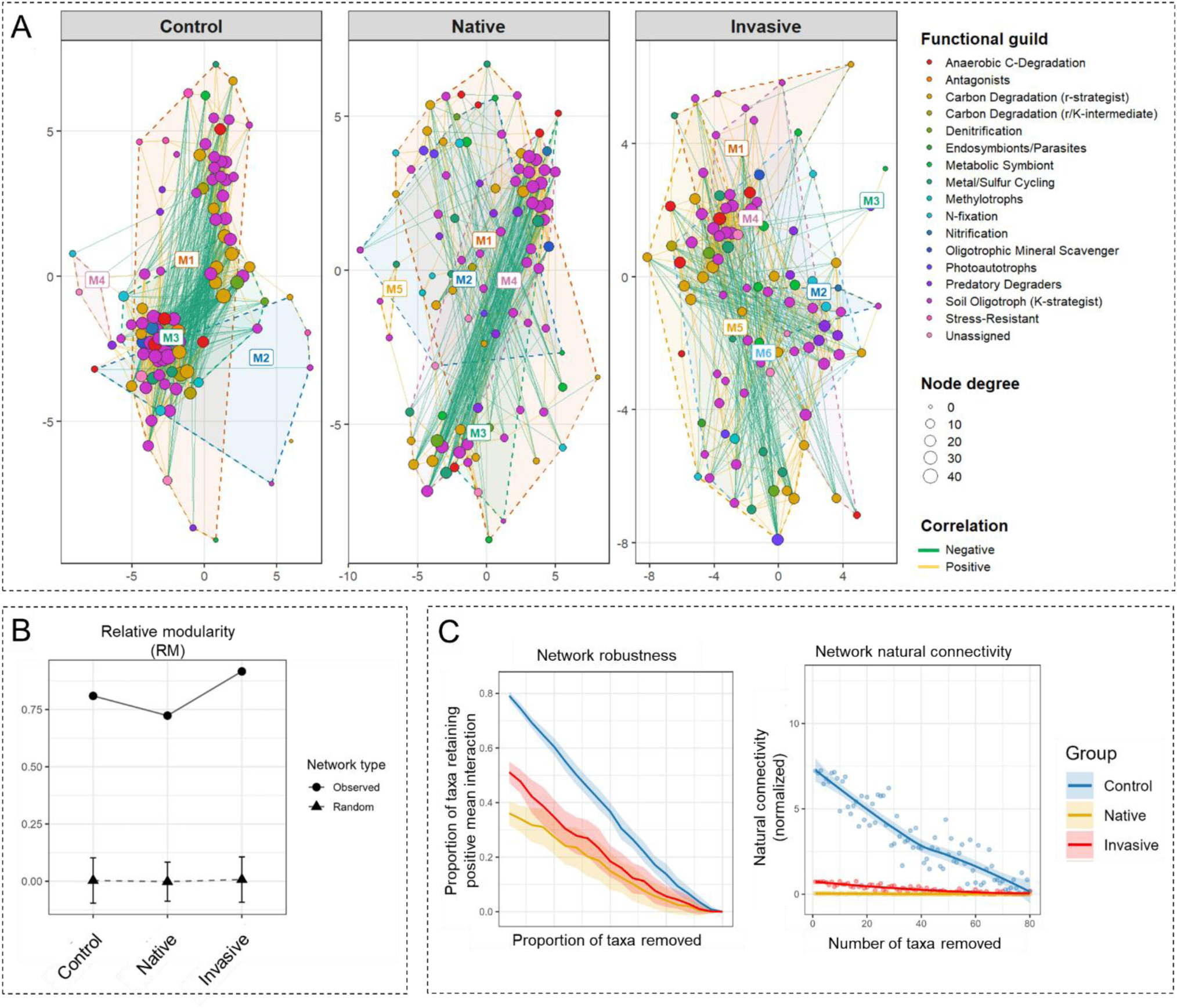
Soil legacy effects on functional-guild organization and robustness of family-level co-occurrence networks. (A) Family-level co-occurrence networks for control, native-conditioned, and invasive-conditioned soils, generated by pooling all incubation treatments within each soil legacy. Nodes represent microbial families, are colored by functional guild, and are scaled by degree. Positive and negative associations are shown as yellow and green edges, respectively. Dashed outlines indicate network modules. (B) Comparison of observed network modularity with corresponding random networks for the three soil types. (C) Network robustness under simulated taxon removal. Left, proportion of taxa retaining positive mean interaction as an increasing proportion of taxa is removed. Right, normalized natural connectivity as taxa are sequentially removed. Shaded ribbons indicate variation among simulations. Overall, the figure shows that soil legacy altered both the internal organization of guild-affiliated families and the robustness of the resulting family-level association networks.

The distribution of selected genera derived from the core microbiome, together with taxa identified by LEfSe across all soils and by ALDEx2 in native-versus-invasive comparisons, was examined (Fig. 5). Several selected genera were highly prevalent across samples, indicating that control, native-, and invasive-conditioned soils shared a substantial common taxonomic backbone (Fig. 5A), where the relative-abundance distribution of these selected genera varied mainly among soils (Fig. 5B). Thus, differences among soils were not expressed as a major replacement of dominant taxa, but rather as shifts in the relative contribution of selected shared and differential genera within a broader background assemblage. This pattern suggested that soil legacy and short-term amendments influenced community organization more through redistribution of shared taxa than through strong taxonomic turnover. Family-level co-occurrence networks were then constructed to determine whether these compositional shifts were accompanied by changes in interaction structure and robustness (Fig. 6).

### 3.4 Soil legacy effects on functional-guild organization and robustness of family-level co-occurrence networks

Family-level co-occurrence networks were used to test whether the distribution of shared taxa described above was accompanied by changes in interaction structure and stability (Fig. 6). All three soil networks retained a similar number of families, supporting that control, native-, and invasive-conditioned soils shared a broad family-level backbone. However, their topology differed clearly (Fig. 6A). The control network was the most connected overall, comprising 99 families and 978 edges, with the highest connectance (0.202), mean degree (19.76), clustering coefficient (0.564), and centralization values. Both plant-conditioned soils contained 100 families but fewer edges and lower overall connectivity than the control, indicating that plant conditioning reduced the density of family-level associations relative to unplanted soil. The invasive network was more connected and more compact than the native network. It contained more edges than the native network (636 vs. 522), a greater proportion of positive associations (77.2% vs. 65.9%), higher connectance (0.128 vs. 0.105), and a higher mean degree (12.72 vs. 10.44), while also showing a shorter average path length (1.57 vs. 1.81). By contrast, the native network was sparser and contained relatively more negative associations. Clustering coefficients were similar in native and invasive soils (0.433 and 0.428, respectively), but both were lower than in the control network.

The three soils also differed in module structure, with four modules in the control network, five in the native network, and six in the invasive network. This progressive increase in module number suggests repartitioning of shared families rather than community-members turnover. Comparison with random networks further showed that observed modularity exceeded random expectations across the three soils (Fig. 6B), indicating non-random compartmentalization of family-level associations. Robustness analysis showed a complementary pattern (Fig. 6C). Under simulated taxon removal, the control network retained the highest proportion of taxa with positive mean interactions across the removal gradient, indicating the greatest overall robustness. Both plant-conditioned networks were less robust than the control, but the invasive network generally retained more structure than the native network. In the weighted analysis, after removal of 5% of taxa, the remaining proportion was 0.791 in the control, 0.511 in the invasive network, and 0.360 in the native network; after removal of 50% of taxa, the corresponding values were 0.367, 0.185, and 0.149, respectively. The same overall ranking was observed in the unweighted analysis, and natural connectivity likewise remained highest in the control and generally intermediate in the invasive network relative to the native network (Supplementary Figures S1 and S2). These results show that soil legacy influenced family-level network organization mainly by changing how shared families were connected, partitioned, and retained under perturbation. Relative to the native-conditioned soil, the invasive-conditioned soil supported a network that was denser, more positive-association dominated, and more compact, whereas both plant-conditioned soils remained less robust than the unplanted control.

To summarize the native-to-invasive rewiring pattern, we next synthesized the dominant module transitions and topological-role shifts between the two plant-conditioned networks (Fig. 7, Supplementary results). The shift from native- to invasive-conditioned soil was not random, but was associated mainly with taxa assigned to the Soil Oligotroph (K-strategist) and Carbon Degradation (r-strategist) guilds. Many of these taxa changed topological position primarily through exchanges between peripheral and connector roles, indicating redistribution of between-module linkage rather than simple gain or loss of taxa. Within the oligotroph guild, some families that acted as connectors in the native network, such as UBA12225 and Gp7-AA8, became peripherals in the invasive network, whereas others, including Burkholderiales_WYBJ01, QHBQ01_JADMIH01, and UBA2386, shifted in the opposite direction and became connectors under invasive conditioning. A comparable pattern was observed for carbon-degradation guilds: *Chitinophagaceae*, *Propionibacteriaceae*, and *Hymenobacteraceae* gained connector roles in the invasive network, whereas *Microbacteriaceae* showed the reverse shift. Thus, the native-to-invasive transition was characterized not by uniform centralization of all taxa, but by selective reassignment of module-linking roles among guilds involved in conservative soil persistence and carbon processing. This pattern suggests that invasion legacy altered how shared taxa were positioned across the network, potentially changing the routing of labile-carbon processing and cross-module associations. Because root exudates are a major source of soluble carbon and a likely driver of rhizosphere priming, these rewiring patterns motivated Experiment 2, in which we directly tested whether native and invasive exudates differentially express the legacy-dependent functional organization inferred from the network analysis.

**Figure 7.**
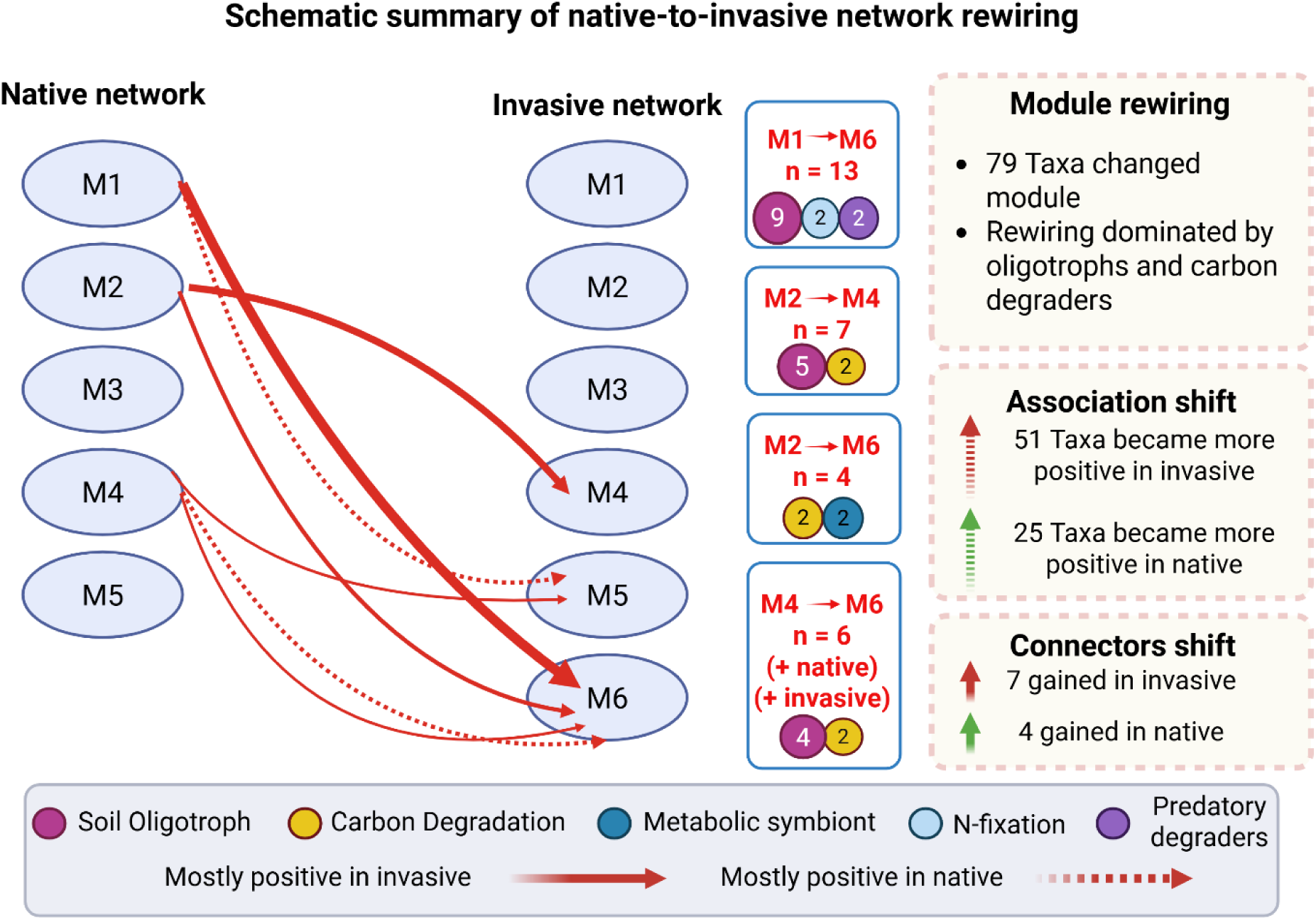
Conceptual illustration of native-to-invasive network rewiring. Major module shifts between native- and invasive-conditioned family-level co-occurrence networks are shown schematically. Circles represent network modules, arrows indicate dominant native-to-invasive transitions, arrow width reflects the number of taxa involved, and line style indicates whether shifted taxa were more positive in the invasive (solid) or native (dashed) network. Colored symbols identify the principal functional guilds contributing to each transition. Summary boxes highlight the extent of module rewiring, the overall direction of association shifts, and connector changes between the two soil legacies. Created in BioRender.

### 3.5 Root exudates elicit soil-legacy-dependent functional responses in conditioned soils (Experiment 2)

Experiment 1 established that soil legacy shaped microbial function and association structure under simplified resource pulses, as reflected in the functional response (Fig. 2) and network differences summarized in Fig. 6. Experiment 2 built on this result by asking whether those same legacy effects would be expressed under more biologically relevant inputs, namely root exudates from the native and invasive plants themselves. By contrasting native and invasive exudates with glucose, Experiment 2 allowed to distinguish general responses to labile carbon from soil-legacy-dependent responses to plant-specific chemical inputs, thereby providing a more mechanistic interpretation of the functional differentiation observed in Experiment 1. Experiment 2 showed that short-term responses to exudate addition depended on both prior soil conditioning and exudate identity, and that invasive and native exudates did not elicit equivalent responses (Fig. 8). Across variables, invasive exudates generally produced clearer separation among soils than native exudates, particularly for the enzyme-based response variables. For β-1,4-N-acetylglucosaminidase activity (Fig. 8A), native exudates did not significantly differentiate the three soils, whereas invasive exudates did. Under invasive exudate addition, activity was higher in control soil than in native-conditioned soil, with invasive-conditioned soil remaining intermediate. This indicates that the invasive exudate generated a sharper short-term expression of soil legacy than the native exudate for this N-acquiring enzyme. An even stronger contrast was observed for α-D-glucosidase activity (Fig. 8B). Invasive exudates produced full separation among all three soils, with the highest activity in control soil, intermediate values in invasive-conditioned soil, and the lowest activity in native-conditioned soil. In contrast, native exudates did not produce significant differences among soils. Thus, for both extracellular enzymes, invasive exudates elicited a stronger and more discriminating functional response than native exudates over the 20 h incubation. nirS abundance showed a more moderate but still legacy-dependent response (Fig. 8C). Under both invasive and native exudate additions, control soil exceeded native-conditioned soil, while invasive-conditioned soil remained intermediate. This suggests that nirS-carrying communities responded to exudate addition in a soil-dependent manner, but unlike the enzyme activities, the contrast between native and invasive exudates was less pronounced. In contrast, archaeal 16S rRNA gene abundance did not differ significantly among soils under either exudate treatment (Fig. 8D), indicating that archaeal abundance was comparatively insensitive to exudate identity over this short timescale. Any short-term response was therefore weaker at the archaeal DNA level than for the functional markers. Overall, these results show that plant exudates did not act simply as interchangeable labile inputs. Rather, exudate identity shaped how preconditioned soils expressed microbial responses, with invasive exudates producing broader and stronger short-term differentiation among soil legacies than native exudates, especially for extracellular enzyme activities.

**Figure 8.**
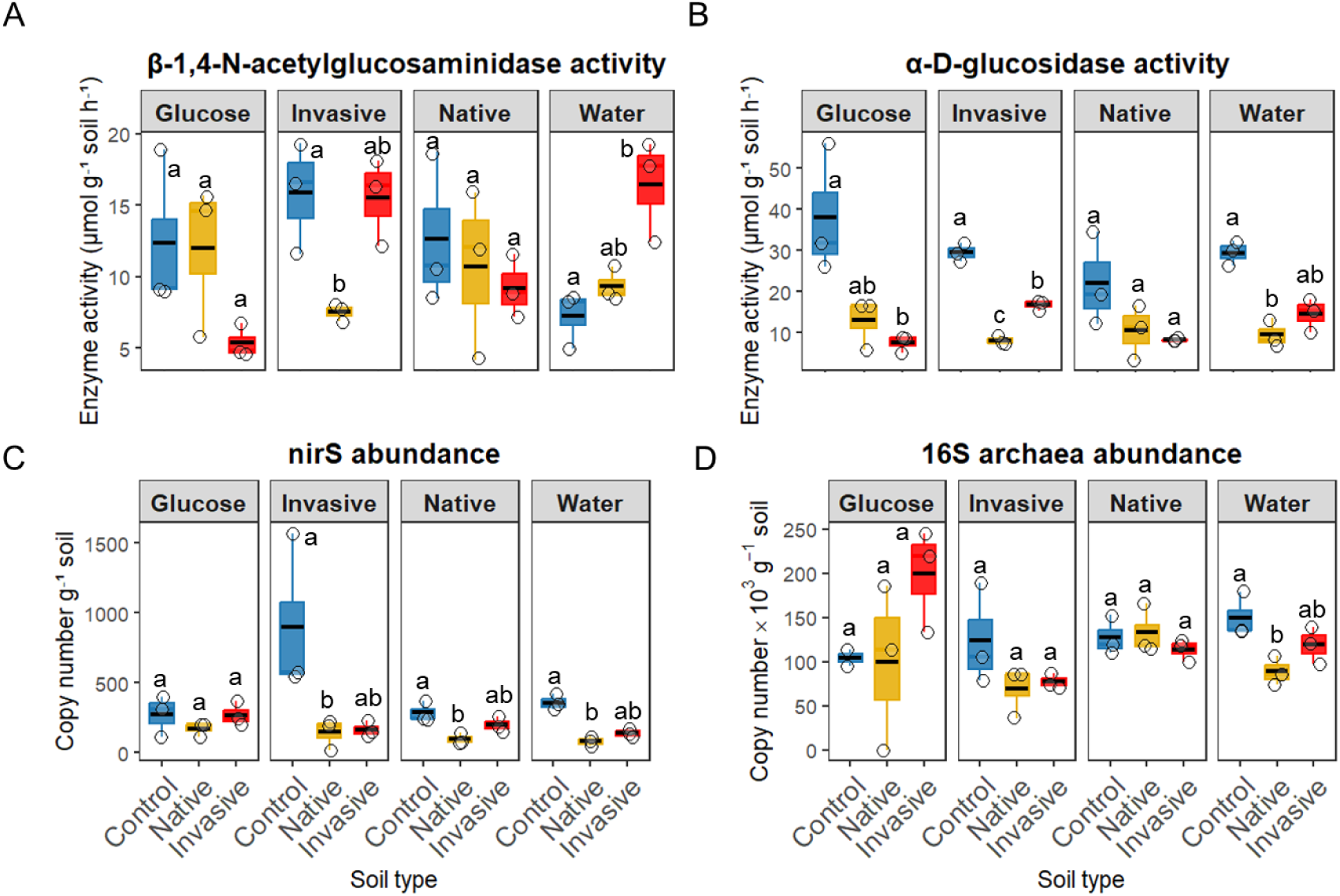
Soil-legacy-dependent microbial responses to short-term additions of glucose and plant root exudates in Experiment 2. Responses of β-1,4-N-acetylglucosaminidase activity, α-D-glucosidase activity, nirS abundance, and archaeal 16S rRNA gene abundance after 20 h incubation of control, native-conditioned, and invasive-conditioned soils amended with glucose, invasive root exudates, native root exudates, or water. Root exudates were collected hydroponically from sterile-grown native (*Helminthotheca echioides*) and invasive (*Conyza bonariensis*) plants. Boxplots show medians and interquartile ranges, and points represent biological replicates; technical replicates were averaged before analysis. Letters indicate significant differences among soil types within each amendment treatment, based on separate ART ANOVAs followed by Tukey-adjusted post hoc tests. Overall, invasive exudates produced stronger short-term soil-legacy differentiation than native exudates, particularly for extracellular enzyme activities and nirS abundance, whereas archaeal 16S abundance responded weakly.

## 4. Discussion

Our results indicate that plant conditioning altered belowground processes mainly by reshaping the functional organization and interaction structure of a broadly shared microbial assemblage, rather than by causing strong taxonomic replacement. In Experiment 1, native- and invasive-conditioned soils differed significantly in composition, yet they were not consistently distinguished by strong family-level indicator taxa. This pattern was reinforced by the network analysis, in which native and invasive soils showed similar node numbers but differed in connectivity, modularity, and robustness. Collectively, these results suggest that plant legacy was expressed primarily through shifts in community organization and interaction architecture, rather than through wholesale turnover of dominant taxa. This interpretation is consistent with invasion literature showing that microbial responses to invasive plants are often heterogeneous and context dependent, and that compositional and functional shifts do not always coincide with major changes in alpha diversity. It also fits plant–soil feedback theory, which predicts that legacy effects accumulate through numerous subtle modifications in soil biota and soil properties over time, rather than through abrupt community replacement (Custer and van Diepen, 2020; Elgersma et al., 2011; Malacrinò et al., 2020; Torres et al., 2021).

A central outcome of both experiments was that functional responses were clearer and faster than broad taxonomic responses. In Experiment 1, short-term cellulose and ammonium additions altered NAGase activity and nirS abundance, but alpha diversity remained driven more strongly by incubation condition than by differences between native- and invasive-conditioned soils. In Experiment 2, this contrast became even clearer: extracellular enzyme activities responded more strongly and more consistently to glucose and root exudates than archaeal abundance. This supports the view that short-term resource pulses can first be expressed through changes in microbial activity, substrate use, and functional potential before they become visible as stronger shifts in community membership. It also aligns with current understanding that root exudates act not only as carbon and nutrient sources, but also as biochemical signals that shape rhizosphere assembly and plant–soil feedback outcomes. Together, our data suggest that invasion legacy persists as a functional filter that conditions how the resident microbiome responds to both simple substrates and biologically relevant plant-derived inputs.

The network results provide an important extension of this interpretation by showing that the native and invasive legacies differed not simply in composition, but in the organization of inferred associations among shared families. The invasive-conditioned soil supported a denser, more positively connected, more compact, and more strongly modular network than the native-conditioned soil, while both plant-conditioned networks were less robust than the unplanted control. This suggests that invasion legacy reorganized the microbial community into a more tightly structured association network rather than replacing its dominant members. Similar studies have shown that network properties can reveal ecological complexity not captured by composition alone, including changes in modular structure and resilience under disturbance. At the same time, co-occurrence networks should be interpreted cautiously: they represent inferred associations rather than direct ecological interactions, and some edges can be environmentally driven rather than biotically mediated. Thus, the present network analysis is best viewed as evidence for rewiring of community organization, not proof of direct pairwise interactions.

The rewiring signal itself was ecologically informative. Dominant transitions between native and invasive modules were driven mainly by soil oligotrophs and carbon-degrading groups, with most module-shifting taxa showing a more positive signed balance in the invasive network. That pattern is consistent with the idea that invasive conditioning can alter not only who is present, but how microbial guilds are arranged across resource-use space. In our system, oligotroph- and carbon-degrader-dominated shifts suggest that invasion legacy changed the balance between conservative resource scavenging and labile-carbon processing functions within the shared family pool. The response to exudates in Experiment 2 reinforces that interpretation. Because root exudates are chemically diverse and under plant control, they can recruit, stimulate, or suppress distinct subsets of the rhizosphere microbiome, thereby altering microbial functional trajectories and subsequent plant–soil feedbacks. Our results suggest that such filtering may persist even after conditioning, so that amended soils differ less in the presence of completely unique taxa than in how shared taxa are activated and reorganized.

An additional insight from this study is that the unplanted control was not simply an intermediate state, but represented a distinct structural benchmark. The control soil showed the highest connectivity and robustness, while both plant-conditioned soils were less robust under simulated taxon removal. This could indicate that plant conditioning, whether by native or invasive species, imposes selective structuring that increases compartmentalization but reduces overall redundancy relative to the unplanted background soil. However, the control must be interpreted with caution in this study because it was not incubated and watered alongside the planted soils during the conditioning phase. For that reason, the contrast between control and planted soils should not be attributed solely to plant identity. Even so, the contrast between native and invasive soils remains informative, because those two legacies were generated under the same conditioning framework and therefore isolate plant-history effects more directly.

From an applied soil ecology perspective, the combined results suggest that microbial legacy effects may be most detectable through function-sensitive and organization-sensitive metrics, rather than through alpha and beta diversity indices alone. In practical terms, extracellular enzyme activities, targeted N-cycle markers, and network-level descriptors may offer earlier or more informative indicators of invasion legacy than taxonomic richness by itself. This is relevant for restoration and management, where soils may appear compositionally similar yet remain functionally and structurally distinct because of persistent legacy effects. More broadly, recent work in both natural and agricultural systems has emphasized that plant-derived soil legacies and root exudate chemistry can be harnessed or redirected to alter microbiome trajectories. Our findings support that applied view, but also indicate that legacy effects can persist across contrasting short-term inputs, making them important to consider when designing restoration, revegetation, or microbiome-informed soil management strategies.

## 5. Conclusions

Plant conditioning by native and invasive species left a measurable microbial legacy, but that legacy was expressed mainly through functional differentiation and network rewiring of a broadly shared microbiome, rather than through strong taxonomic turnover. Across both experiments, native- and invasive-conditioned soils differed in enzyme activity, nirS responses, and association structure, even when differences in alpha diversity or dominant indicator taxa were limited. The invasive-conditioned soil supported a denser and more positive network than the native-conditioned soil, and the native-to-invasive transition was driven largely by shifts in peripheral versus connector roles among oligotrophic and carbon-degrading guilds. Experiment 2 showed that these legacy effects were not restricted to simplified amendments, but were also expressed under plant-derived inputs, with invasive exudates producing stronger short-term functional differentiation than native exudates. Together, these findings indicate that invasion can leave a persistent belowground signature that alters how resident microbiomes process subsequent inputs, with implications for plant–soil feedbacks, carbon turnover, and soil management.

## Supporting information

Supplementary Results

## Acknowledgements

We gratefully acknowledge the Israel Ministry of Innovation, Science and Technology for funding project number 4673 (Investigating greenhouse gases emissions in plant invasion hotspots as a model for aquatic ecosystem restoration and management), which supported the research efforts leading to the development and writing of this paper.

Data availability

