## Supplementary Methods for "Invasive plant soil legacies shape microbial function and community organization under short-term carbon and nitrogen amendments"

### 1. Composition and preparation of Hoagland's medium used for root exudate collection

For root exudate collection, plants previously established in Magenta boxes were transferred to Hoagland's nutrient solution prepared from concentrated stock solutions. Macronutrient stock solutions were prepared separately in 50 mL distilled water as described in Table S1.

Table S1. Composition of Hoagland's medium used for root exudate collection

| Component | Stock composition |
|---|---|
| **$Ca(NO_3)_2 \cdot 4H_2O$ stock** | 11.8 g in 50 mL distilled water |
| **$KNO_3$ stock** | 5.1 g in 50 mL distilled water |
| **$KH_2PO_4$ stock** | 6.8 g in 50 mL distilled water |
| **$MgSO_4 \cdot 7H_2O$ stock** | 12.3 g in 50 mL distilled water |
| **Micronutrient stock (1000×)** | $H_3BO_3$, 0.14 g; $MnCl_2 \cdot 4H_2O$, 0.09 g; $ZnSO_4 \cdot 7H_2O$, 0.01 g; $CuSO_4 \cdot 5H_2O$, 0.001 g; $Na_2MoO_4 \cdot H_2O$, 0.001 g, in 50 mL distilled water |
| **Fe-EDTA stock** | EDTA, 10.4 g; $FeSO_4 \cdot 7H_2O$, 7.8 g; KOH, 56.1 g, brought to 50 mL total volume. The pH of this stock was adjusted to 5.5 using $H_2SO_4$, |

The pH of the final medium was adjusted to 5.8–6.0.

Working solution (1×, per 50 mL final volume): 2.5 mL $Ca(NO_3)_2 \cdot 4H_2O$ stock, 2.5 mL $KNO_3$ stock, 0.5 mL $KH_2PO_4$ stock, 1.0 mL $MgSO_4 \cdot 7H_2O$ stock, 0.5 mL micronutrient stock, and 0.5 mL Fe-EDTA stock; final pH adjusted to 5.8–6.0

### 2. Extracellular enzyme assays

Potential activities of two common hydrolytic enzymes were measured to evaluate microbial capacity for degradation of soil carbohydrate and nitrogen-containing organic substrates: α-1,4-glucosidase (AG; EC 3.2.1.20), which hydrolyzes terminal α-glucosidic linkages in starch-like compounds, and β-1,4-N-acetylglucosaminidase (NAGase; EC 3.2.1.52), which cleaves N-acetyl-β-D-glucosamine from chitin- and peptidoglycan-derived substrates. Assays followed modified fluorometric protocols based on German et al. (2011) and Hless and Yanuka-Golub (in press), using 4-methylumbelliferyl α-D-glucopyranoside (Sigma-Aldrich 69591) for AG and 4-methylumbelliferyl N-acetyl-β-D-glucosaminide (Sigma-Aldrich M2133) for NAGase. Standard curves were prepared using 4-methylumbelliferone sodium salt dissolved in sterile deionized water to a stock concentration of 600 µM and serially diluted to generate working standards. For final activity calculations, standard concentrations were corrected to the molar mass of neutral MUF, corresponding to the fluorophore released during enzymatic hydrolysis.

Soil homogenates were prepared fresh on the day of analysis by vortexing 2.0 g dry-weight-equivalent soil in 10 mL sterile deionized water for 10 min at half speed (Vortex Genie 2, Scientific Industries). Each assay included three technical replicates per sample and substrate concentration, together with the following controls: homogenate control (soil plus water without substrate), substrate control (substrate plus water without soil), homogenate quality control (homogenate spiked with known MUF concentration), MUF standard control, and water blank. All reactions were conducted in a total volume of 2 mL, consisting of 1 mL homogenate or water and 1 mL substrate solution, and were incubated in the dark at room temperature for 1 h.

After incubation, samples were centrifuged at 14,000 rpm for 5 min. An aliquot of 250 µL supernatant was transferred to a black 96-well microplate, and 50 µL sterile water was added to each well. Fluorescence was measured using a Feyond-A300 microplate reader with excitation at 365 nm and emission at 450-460 nm. Fluorescence values were converted to product concentrations using corrected MUF standard curves, with sample-specific quench correction applied from the homogenate-spiked controls. Final enzyme activities were expressed as µmol MUF released $g^{-1}$ dry soil $h^{-1}$ after correction for reaction volume, incubation time, dry soil mass, and background fluorescence.

**3. DNA extraction, PCR amplification, and amplicon sequencing**

Whole-community genomic DNA was extracted from soil subsamples (374 ± 37 mg) using the FastDNA™ SPIN Kit for Soil, according to the manufacturer's instructions with minor modifications, and was stored at −20 °C until further analysis. DNA concentration and purity were assessed using a NanoDrop ND-1000 spectrophotometer (Thermo Scientific, Wilmington, DE, USA). PCR products were verified by electrophoresis on 1.5% agarose gels stained with Hy-View Nucleic Acid Stain (Cat. No. IMGS7011). Amplicon libraries were prepared for sequencing on a PacBio Revio platform using a two-stage PCR approach as described previously (Verma et al., 2025), similar in principle to two-step amplicon library preparation methods used for Illumina sequencing (e.g., Naqib et al., 2018). In the first PCR stage, full-length bacterial 16S rRNA genes were amplified using the domain-level primers 27F/1492R (Gao et al., 2024). First-stage PCR reactions were performed in 30 µL volumes using 2× MyTaq HS Mix, with the following cycling conditions: initial denaturation at 95 °C for 5 min; 29 cycles of 95 °C for 30 s, 50 °C for 30 s, and 72 °C for 60 s; and a final extension at 72 °C for 5 min.

Second-stage PCR amplification and initial library preparation were performed at the Genomics and Microbiome Core Facility (GMCF) at Rush University. Reactions were carried out in 10 µL volumes in 96-well plates using repliQa HiFi ToughMix (2× mastermix). Each well received a unique primer pair (final concentration 1 µM) containing PacBio Kinnex adapter sequences, combinatorial dual indices, and ONT linker sequences at the 3′ ends. One microliter of unpurified first-stage PCR product was used directly as template for the second-stage reaction. Cycling conditions were 98 °C for 2 min, followed by 8 cycles of 98 °C for 10 s, 60 °C for 1 s, and 68 °C for 2 s. Amplified products were pooled and purified using a 0.5× AMPure cleanup. Final Kinnex library preparation and PacBio sequencing were carried out at the DNA Services Facility at the Roy J. Carver Biotechnology Center, University of Illinois Urbana-Champaign.

**4. 16S data filtering, normalization, and ordination workflow for Experiment 1.**

The starting ASV table contained 13,934 ASVs and 43 samples, together with five taxonomy columns (Phylum, Class, Order, Family, and Genus). Samples labeled as "Competition" were excluded first, leaving 34 samples. The count matrix was then restricted to those retained samples, and ASVs with zero total abundance across them were removed, resulting in a filtered matrix of 12,548 ASVs × 34 samples. Total read counts were then inspected, and three low-read samples (ContCell3, Cont2_1, and Cont2_2) were removed. After a second zero-abundance filter, the final dataset comprised 12,308 ASVs across 31 samples, with a matched taxonomy table of 12,308 rows × 5 columns.

For alpha diversity, the final ASV matrix was rarefied to 30,000 reads per sample using rrarefy in vegan. Observed richness, Shannon diversity, and Simpson diversity were calculated from the rarefied counts, and effective Shannon and effective Simpson diversity were expressed as Hill numbers. Group differences were tested with one-way ART ANOVA models in ARTool, followed by Tukey-adjusted ART contrasts for factors with more than two levels. A second alpha-diversity analysis was performed on the subset containing only plant-conditioned soils.

Alpha diversity was quantified from the ASV table after filtering samples and ASVs to match the experimental design. Samples labeled as "Competition" were excluded, and three low-read samples (ContCell3, Cont2_1, Cont2_2) were removed prior to analysis. The ASV count matrix was then restricted to the retained samples, and ASVs with zero total abundance across the retained samples were discarded . To account for differences in sequencing depth among the remaining 31 samples, the count matrix was rarefied to a constant depth of 30,000 reads per sample using random subsampling without replacement (function rrarefy in vegan ). From the rarefied counts, we calculated observed richness (number of ASVs), Shannon diversity, and Simpson diversity (vegan::specnumber and vegan::diversity). In addition, we derived effective diversities (Hill numbers) to express diversity in units of "effective number of taxa": effective Shannon diversity as exp(Shannon) and effective Simpson diversity as 1/(1 − Simpson).

Group differences in alpha diversity were tested using aligned rank transform (ART) ANOVA (art function in the ARTool ), which enables factorial ANOVA-style inference on non-normally distributed responses by aligning and ranking the response prior to fitting linear models For each response metric, we tested main effects of Soil (Control, Native, Invasive), Condition (T0, Water, Ammonium, Cellulose), and Plant (Yes/No) in separate one-way ART models. Full factorial models including soil and condition interactions were not used because the per-cell replication was low (typically 2–3 samples per Soil × Condition combination), limiting stable estimation and interpretation of interaction effects. When a factor with more than two levels was significant, post hoc pairwise contrasts were computed using ART-based comparisons (ARTool::art.con ) with Tukey adjustment for multiple testing. A second analysis was conducted on the subset of samples containing plants to directly compare Native versus Invasive soils without the Control soils.

For community composition, the non-rarefied ASV table was normalized by CSS using newMRexperiment, cumNormStatFast, and cumNorm in metagenomeSeq. The resulting CSS-normalized ASV matrix retained dimensions of 12,308 × 31. Counts were then summed by taxonomic assignment to the genus level, yielding a 1,271 × 31 genus matrix. Bray–Curtis dissimilarities were calculated from both ASV- and genus-level CSS-normalized matrices and visualized by two-dimensional NMDS (metaMDS, k = 2, distance = "bray", try = 100, trymax = 100, autotransform = FALSE). The final stress values were 0.03593292 for the ASV-level ordination and 0.03861967 for the genus-level ordination. A second plant-only genus-level ordination was generated after exclusion of the control soil, and its stress value was 0.1537943.

Community-composition differences were tested by PERMANOVA using adonis2 on Bray–Curtis distances. In the full dataset, significant effects were detected for Soil (genus level: $R^2$ = 0.449, F = 41.59, P = 0.001), Condition ($R^2$ = 0.229, F = 14.15, P = 0.001), and Soil × Condition ($R^2$ = 0.219, F = 6.75, P = 0.001). At the ASV level, the same factors were also significant, although with lower explained variance for Soil ($R^2$ = 0.344), Condition ($R^2$ = 0.156), and Soil × Condition ($R^2$ = 0.203). Pairwise PERMANOVA tests were additionally performed for Soil and Condition contrasts. For the plant-only subset at genus level, significant effects were detected for Soil ($R^2$ = 0.0937, F = 4.14, P = 0.001), Condition ($R^2$ = 0.4565, F = 6.72,

P = 0.001), and Soil × Condition ($R^2$ = 0.1329, F = 1.96, P = 0.013). Pairwise contrasts among conditions showed significant differences for Water vs Ammonium (P = 0.002), Water vs T0 (P = 0.002), Ammonium vs Cellulose (P = 0.004), Ammonium vs T0 (P = 0.003), and Cellulose vs T0 (P = 0.009), whereas Water vs Cellulose was marginal (P = 0.059).

**5. Differential abundance analyses**

Differential abundance analyses were performed on the plant-conditioned dataset using both LEfSe and ALDEx2. LEfSe was run with Soil as the grouping variable, without subgroup stratification, using either ASV-level features or all taxonomic ranks. Analyses were performed with identity transformation and no additional normalization, with Kruskal–Wallis cutoff = 0.99, Wilcoxon cutoff = 0.05, LDA cutoff = 0.01, 30 bootstrap iterations, bootstrap fraction = 2/3, and minimum subclass sample size = 10. Marker tables were exported for downstream interpretation. In parallel, condition-specific pairwise differential abundance analyses between native- and invasive-conditioned soils were performed at the genus level using ALDEx2. For each incubation condition (Water, Ammonium, and Cellulose), taxa present in at least three samples were retained, CSS-normalized counts were rounded to integers, and soil legacy was used as the grouping factor. ALDEx2 was run using the Kruskal–Wallis test with effect-size estimation and the denominator set to all features. To aid interpretation, mean abundances were calculated for each soil × condition combination and fold-change values between invasive- and native-conditioned soils were extracted alongside unadjusted and Benjamini–Hochberg-adjusted P values.

**6. Network structure and stability**

Data preparation

ASV data from Experiment 1 were aggregated to the family level and normalized using cumulative sum scaling (CSS) implemented in the metagenomeSeq R package (v1.52.0). Co-occurrence networks and downstream analyses were conducted using ggClusterNet (v2.0). All analyses were performed in parallel for two experimental factors, soil treatment (Control, Native, Invasive) and incubation treatment (T0, Water, Ammonium, Cellulose), yielding seven group-specific networks. Interaction networks for specific soil × incubation combinations were not constructed because per-combination sample sizes were insufficient for stable co-occurrence inference.

Network construction

Networks were inferred with ggClusterNet::network.pip using Spearman rank correlations computed on CSS-normalized family abundances. Within each factor level, the top N = 100 families were selected based on total CSS abundance within that level and used for network construction. Pairwise associations were retained if $|r| \geq 0.6$ and $p \leq 0.05$, and the resulting adjacency matrices were used as the group-specific networks for downstream analyses.

Node-level analyses

To evaluate whether observed connectivity patterns differed from random expectation, degree distributions were compared against Erdős–Rényi (E–R) random networks generated within ggClusterNet (ram.net = TRUE, R = 5). Node degree was defined as the number of edges incident to a node. For each group-specific network, the proportion of taxa at each degree value was computed for the observed network and the corresponding random networks. Node topological roles were quantified using within-module connectivity (Zi) and participation coefficient (Pi) following Guimerà and Amaral (2005). Zi measures the strength of within-module connectivity, whereas Pi measures how evenly links are distributed across modules. Nodes were classified using the four-role scheme with thresholds Zi = 2.5 and Pi = 0.62 to distinguish peripherals (Zi < 2.5, Pi < 0.62), connectors (Zi < 2.5, Pi ≥ 0.62), module hubs (Zi ≥ 2.5, Pi < 0.62), and network hubs (Zi ≥ 2.5, Pi ≥ 0.62).

Community structure

Community structure was characterized using modularity-based module detection, and module composition was summarized based on the inferred modules. Modules represent sets of taxa that are more densely connected to each other than to the rest of the network. For each group-specific network, family-level abundances were summarized within modules, reporting both absolute abundances and relative abundances (within-module proportions) for the most abundant families.

Network-level properties

Network structure was summarized using standard graph-level properties extracted from ggClusterNet outputs. For metrics with random-network references, observed values were compared against matched E–R random networks generated within ggClusterNet. Degree and connectance were not interpreted relative to the E–R baseline because the random networks preserve the observed numbers of nodes and edges. The following metrics were computed for each group-specific network:

• Connectance (edge density): proportion of realized edges out of all possible edges among nodes, where higher values indicate a denser network.

• Average degree: mean number of edges per node, where higher values indicate taxa have more associations on average.

• Average path length: mean shortest-path distance between pairs of nodes, where lower values indicate a more compact and globally connected network.

• Mean clustering coefficient: average tendency for nodes to form triangles, where higher values indicate stronger local clustering.

• Degree centralization: concentration of degree in a few hubs, where higher values indicate stronger hub dominance.

• Betweenness centralization: concentration of shortest-path brokerage in a few nodes, where higher values indicate greater control by potential bottlenecks.

• Closeness centralization: concentration of global reach in a few nodes, where higher values indicate a few nodes are much more centrally located than others.

• Edge connectivity: minimum number of edges that must be removed to disconnect the network, where higher values indicate more robust connectivity.

• Modularity: strength of community structure, where higher values indicate stronger subdivision into modules.

• Proportion of negative edges: percentage of inferred associations with negative correlation, where higher values indicate a greater prevalence of negative associations.

Network stability

Network robustness to random taxon loss was evaluated using ggClusterNet:::Robustness.Random.removal. For each group-specific network, taxa were progressively removed at random and robustness was summarized as the proportion of taxa retaining positive mean interaction strength in the reduced network. Robustness was assessed for both unweighted networks and abundance-weighted networks, and results were summarized across permutations for each removal level. Network stability was additionally assessed using natural connectivity implemented with ggClusterNet:::natural.con.microp. Natural connectivity quantifies the redundancy of alternative paths in a network, with higher values indicating a more resilient structure. For each group-specific network, increasing numbers of taxa were removed at random (0–80 removals) and natural connectivity was computed for the remaining subnetwork.
